# Gene expression response to temperature acclimation in a model photoheterotrophic marine flavobacterium (*Dokdonia* sp. MED134)

**DOI:** 10.1101/2023.12.25.573297

**Authors:** Semidán Robaina-Estévez, José M. González, Ana María Cabello, Antonio S. Palacio, Ruairí Gallagher, Ángel López-Urrutia, Laura Alonso-Sáez

## Abstract

Temperature stands as one of the key factors influencing the community structure and distribution of marine microorganisms. Yet, the physiological adaptations of marine bacteria to rising temperatures remain underexplored. This study examines the transcriptional response of *Dokdonia* sp. MED134, a proteorhodopsin-based phototrophic organism within the class *Flavobacteriia*, to a gradient of temperature acclimation conditions from 10 to 34°C. Light availability during day/night cycles exerted minimal influence on the transcriptional patterns of this strain, with only a few genes mostly related to light sensing, light protecting mechanisms and phototrophy being upregulated during daytime. By contrast, temperature significantly impacted the expression of a large fraction of MED134 genes (>60%), including components of the stress response, cellular translation, DNA replication, and some metabolic pathways such as the anaplerotic carbon fixation and the glyoxylate shunt, suggesting intracellular carbon flow adjustments to temperature. Notably, the expression of some highly expressed TonB transporters, prominent in flavobacteria, was also temperature-sensitive. Our findings provide insights into the transcriptional adjustments of *Dokdonia* sp. MED134 in response to temperature variations, suggesting potential implications for carbon cycling and organic matter processing in marine environments.

## INTRODUCTION

Temperature is considered one of the main environmental factors impacting oceanic biogeochemical cycles (Sarmiento et al. 2004). Heterotrophic prokaryotes (including bacteria and archaea, hereinafter referred to as bacteria) have a paramount role in turning such cycles, by e.g., mediating the remineralization of massive amounts of dissolved organic carbon produced daily by primary producers (Boyd et al. 1999; Robinson and Williams n.d.) and controlling fluxes of energy in the ocean (Farooq Azam 1998; F. Azam et al. 1994; le B. Williams and J 1998). Thus, a key question is how temperature impacts fundamental aspects of bacterial metabolism in oceanic waters. The analysis of temperature effects on bulk bacterial metabolic rates, including biomass production, respiration, or enzymatic activity has been the focus of a number of studies (see for example White et al., 1991, Vázquez-Domínguez et al. 2007, Ayo et al. 2017). Temperature also impacts individual bacterial phenotypic traits such as cell size (Morán et al. 2015) and even global patterns of marine surface community composition (Sunagawa et al. 2015). Therefore, the impact of temperature on marine bacterial ecology and functioning is multifaceted and far-reaching, with yet unknown consequences in the present climate-changing scenario.

The metabolic response of bacteria to temperature is not uniform, with even co-existing taxa exhibiting different sensitivities (Pittera et al. 2014; Arandia-Gorostidi et al. 2017). Additionally, we still lack basic knowledge about how heterotrophic marine bacteria respond to temperature variations at the molecular level. Most previous studies with model bacteria have focused on their transcriptional response to rapid temperature down- or upshifts, often forcing cells over their physiological thermal thresholds (Roncarati and Scarlato 2017; Zhang and Gross 2021). However, marine microbial communities will face more gradual temperature variations in a future warming scenario, typically falling within a narrower thermal range. Thus, we urgently need to increase our knowledge of the molecular response of ecologically relevant model bacteria to temperature variations within a moderate thermal range.

Here, we focused on a model strain of the genus *Dokdonia*, affiliated with *Flavobacteriia* (phylum *Bacteroidetes*). This class represents a significant fraction of the oceanic bacterioplankton (Cottrell and Kirchman 2000; Glöckner, Fuchs, and Amann 1999; Abell and Bowman 2005; Alonso et al. 2007) and marine aggregates (O’Sullivan et al. 2004; Rath et al. 1998), where they likely play an important role in the remineralization of sinking carbon. *Flavobacteriia* are well-adapted to grow on marine polymeric substances including proteins and polysaccharides (González et al. 2011; Unfried et al. 2018), which are major carbon and energy sources available in algal exudates (see (Mühlenbruch et al. 2018) for a review).

Additionally, *Flavobacteriia*, and in particular members of the genus *Dokdonia*, including the strain MED134, are also models of photoheterotrophy, a widespread lifestyle of surface marine bacterioplankton that combines heterotrophic growth with light-driven energy production via proteorhodopsins (PR). This extra energy source may be used in different ways, from starvation survival to growth enhancement (Pinhassi et al. 2016), or to cope with the stress associated with salt and nutrient deprivation (Feng et al. 2013).

Understanding the thermal response of PR-containing bacteria is highly relevant, given their predominance in surface ocean communities. The ubiquitous PR protein could be involved in modulating the response of marine bacteria to suboptimal growth conditions (Fuhrman, Schwalbach, and Stingl 2008). In this study, by analyzing the transcriptional response of the strain *Dokdonia* sp. MED134 growing along a range of temperatures encompassing their thermal niche (10 to 34ºC), we aimed at understanding the molecular basis of thermal acclimation of this model photoheterotrophic marine flavobacterium. Beyond the canonical cold- and heat-shock molecular response, our focus was to analyze whether the expression of major physiological and metabolic processes with great ecological relevance for oceanic surface bacterioplankton (i.e., photoheterotrophy and central carbon metabolism, including anaplerotic fixation pathways) changed along their thermal niche. Obtaining such fundamental knowledge is crucial to understanding and predicting how the metabolic landscape of marine bacterioplankton will change in a future warmer ocean.

## MATERIALS AND METHODS

### Culture conditions and RNA extraction

*Dokdonia* sp. MED134 strain was obtained from ICM-CSIC and preserved in glycerol (25% final concentration) at -80ºC. The strain was grown on marine agar plates (1.5% agar; Difco) and maintained at 10, 18, 25, and 34ºC under 12:12-hour light/dark cycles at 180 µM photons m^-2^ s^-1^ light intensity. After two days of growth on agar plates, MED134 colonies from each temperature were inoculated into 5 mL marine broth (Difco) in 15 mL tubes (Falcon) and cultivated overnight with agitation (140 rpm) at their respective temperature. Next, 2-5µL of overnight cultures was transferred to 2 × 50 mL vented-cap glass bottles (Pirex) with 20 mL culture medium with artificial seawater (35 PSU; 40g Red Sea Salt per liter) containing L-alanine (Ala) as a carbon source (final concentration 0.7 mM, see Palovaara et al. 2014 for medium preparation details).

Three replicate cultures at each temperature were grown under these experimental conditions for ca. 3 days under a 12:12 light/dark cycle, and then a volume was transferred to 3 × 1 L polycarbonate bottles filled with the same culture medium (500 mL final volume), to obtain an initial concentration of 5 × 10^4^ cells/mL. Subsequently, in this last cell transfer cells were grown until they reached a target concentration of ca. 1 × 10^6^ cells/mL. At this point, cells were growing exponentially and at a sufficient density for obtaining RNA samples. For monitoring cell density during the acclimation experiments we used flow cytometry (BD Accuri C6 Plus), with samples being collected and fixed with paraformaldehyde and glutaraldehyde (1% and 0.05% final concentration, respectively) at different time intervals (typically each 2-4 hours). Additionally, samples were also preserved during the last cell transfer and analysed in a FACScalibur flow cytometer (Becton Dickinson) equipped with a blue (488-nm) argon laser for obtaining accurate estimates of cell size. Estimates of cell diameter were obtained based on the natural logarithmic-transformed side scatter (SSC) using the calibration provided for heterotrophic bacteria in Calvo-Díaz and Morán, 2006 (Calvo-Díaz and Morán 2006) and cell size was calculated assuming a spherical cell shape.

In the last cell transfer, the target cellular abundance for collecting RNA was reached after 8 to 10 hours of incubation at temperatures 18 to 34ºC, and after 28 hours in incubation at 10ºC, as growth rates were the lowest under the latter temperature. As cells were incubated under a 12h:12h light/dark cycle, they were exposed to light conditions at all sampling times. In order to obtain a proxy of gene expression patterns under dark conditions, a set of replicate bottles was fully covered two hours before the estimated time of RNA collection. In this way, we ensured that cells were acclimated to dark conditions by the time of sampling.

For RNA collection, triplicate culture samples for each light and temperature condition were vacuum filtered onto 0.2 µm PES membranes (Millipore) mounted on Nalgene filtration cups. Each replicate was filtered on two separate filters (250 mL each), with filtration time maintained below 3 minutes. Membranes were flash-frozen in liquid N_2_ and stored at -80ºC until extraction. Replicate filters were separately extracted using the RNAqueous total RNA isolation kit (Ambion) following standard procedures. Six internal RNA standards obtained from *Saccharolobus solfataricus* P2 (https://github.com/Robaina/Dokdonia) were individually spiked to the RNA filters prior to the initiation of the RNA extraction. Replicate RNA extracts per sample were pooled together for library preparation, which was done using the TrueSeq stranded mRNA sample preparation kit (Illumina). cDNA libraries were sequenced as 75-bp paired-end reads on an Illumina HiSeq v4 platform (CNAG, Spain).

### Raw data processing and read counts

Pair-end reads in the FastQ files were first preprocessed to remove low-quality sequences, adapters, and contaminating RNA. To this end, AfterQC was employed (Chen et al. 2017) using default settings, to remove low-quality sequences as well as to trim adapters. rRNA sequences were then filtered out by mapping the transcriptome sequence reads with SortMeRNA version 2.0 (Kopylova, Noé, and Touzet 2012) against a custom-made database of rRNA sequences. This database included all predicted rRNA genes that were found in 28 high-quality *Dokdonia* genome databases described in Baltar et al. (2023). The genomes were screened with Infernal version 1.1.4 (Nawrocki and Eddy 2013) using the models specific to rRNA genes. Phage ΦX174 sequences were removed by aligning the resulting sequence reads with BWA version 0.7.17-r1188 (Li and Durbin 2009) against the sequence of the phage. The BWA alignment files were further processed with samtools version 1.12 (Danecek et al. 2021). After preprocessing, pair-end reads were mapped to the *Dokdonia* MED134 genome (accession number CP009301) using Rsubread’s function *align* (Liao, Smyth, and Shi 2019) with default settings (allowing only one mismatch). Read counts were obtained with Rsubread’s function *featureCount* using default settings for pair-end reads.

Additionally, genes with a read count lower than ten in any of the experimental conditions were removed from the dataset. To obtain counts of the added internal standards, we aligned preprocessed FastQ files to the *Saccharolobus solfataricus* P2 reference genome (NCBI reference ID NZ_CP033238.1) following the same pipeline described above. Quantitative estimates of individual transcript abundance (Ta) of *Dokdonia* MED134 protein-coding genes were obtained following (Gifford et al. 2014) using the calculation Ta = (Ts × Sa) / Ss, where Ta corresponds to the estimated number of transcripts of an individual MED134 protein-coding gene; Ts corresponds to the number of reads assigned to the corresponding MED134 protein-coding gene; Sa corresponds to the molecules of internal RNA standards spiked to the RNA sample, and Ss corresponds to the number of reads assigned to *S. solfataricus* internal standards. In this calculation, one of the standards (standard 14) was removed from the analysis because it was consistently recovered in a higher proportion than other standards. For each sample, Ta values were divided by the total cell abundance in the RNA filter (as obtained by flow cytometry) to obtain estimates of transcript abundance per cell. From the total of 25 replicate samples analyzed, two replicates corresponding to the samples with minimum and maximum cell density at the time of sampling (4.94 × 10^5^ and 1.52 × 10^6^ cells mL^-1^, respectively) exhibited an unusually high and low average number of transcripts per cell, respectively, as compared to the other replicates in the same temperature treatment. These two replicates (L10_R1 and D18_R2) were removed for the calculations of average mRNA transcripts per cell in their corresponding temperature treatment. The final number of replicates considered for the calculation (including day and night samples) in the different temperature conditions ranged from 5 to 8.

### Differential expression analysis

Differentially expressed (DE) genes were obtained with DeSeq2 (Love, Huber, and Anders 2014), a tool developed to analyze differential expression directly from count data via shrinkage estimation for dispersions and fold changes. To analyze differential expression between light and dark conditions for each temperature, we employed the Wald test within DeSeq2, using default settings and with a p-value cutoff of 0.01 and a fold-change cutoff value of 0.5 to compensate for the small number of replicates (≤4) in our experiment (Schurch et al. 2016). To obtain the sets of differentially expressed genes across temperatures – under light, dark, and light or dark conditions – we applied the Likelihood Ratio Test (LRT) within DeSeq2 with default settings and a p-value cutoff of 0.01.

### Gene clustering

Gene clusters were obtained with *Clust*, a clustering algorithm designed to maximize the number of co-expressed genes within clusters (Abu-Jamous and Kelly 2018). *Clust* minimizes the variance within each cluster and does not attempt to assign a cluster to every gene in the sample. As a result, *Clust* reduces the number of spurious assignments of genes that are not truly correlated. *Clust* was employed with default parameters except for the cluster tightness parameter, which was set to three since smaller values decreased the resolution of the clusters producing aggregated clusters of genes with upward or downward trends across temperatures. DeSeq2-transformed counts were the input data to *Clust*. In this analysis, we removed genes that were DE between light and dark conditions (a total of 14 genes; Wald test performed in DeSeq2, ɑ = 0.01). For the remaining genes, light and dark replicate samples were grouped into a single pool as input to *Clust*, i.e., treating light and dark conditions as replicates. Finally, we applied a z-score normalization prior to the clustering step.

### Functional annotations of *Dokdonia* MED134 genome

Manual functional annotations were obtained from González et al. (2011) and additional annotations from KEGG (Kanehisa et al. 2021) under organism code *dok*, corresponding to *Dokdonia* MED134. Operons were predicted using Rockhopper (Tjaden 2020) version 2.03 following the alignment of the transcriptome sequence reads to the genome.

### Pathway over-representation analysis

The R library *clusterProfiler* (Wu et al. 2021) was used to run an over-representation analysis (hypergeometric test with Benjamini-Hochberg correction for multiple testing) of KEGG pathways in each of the clusters previously generated. Specifically, the function *compareCluster* was employed, set to *Dokdonia* MED134 KEGG annotations, a p-value threshold of 0.1, and all genes contained in the clusters as “universe” set for the testing.

### Data availability

All code generated in this study is available in the repository https://github.com/Robaina/Dokdonia and has been organized in a Jupyter Notebook, with Python 3 as the default programming environment. DeSeq2-normalized counts and estimates of transcript per cell for the entire MED134 gene set can be found in Supplementary Table S1. Sequencing reads have been deposited in the European Nucleotide Archive under study accession number PRJEB54885.

## RESULTS

### Growth rate and cell size of MED134 along the thermal niche

*Dokdonia* MED134 cultures were acclimated to a range of four temperatures along their thermal niche under a 12:12 light/dark photoperiod to simulate daily variations in light availability. The growth rate of MED134 increased four-fold from the minimum temperature (Tmin, 10ºC; 3 day^-1^) to 25ºC (12 day^-1^), and entered a thermal stress zone thereafter, demonstrated by a slightly lower growth rate at the maximum temperature (34ºC, 10 day^-1^) (Figure 1A). The average size of MED134 cells showed minor variations over the thermal gradient, with cell diameter varying from 0.60 to 0.63 μm in individual replicates and maximum cell size values at intermediate growth temperatures (Figure 1B).

**Figure 1.**
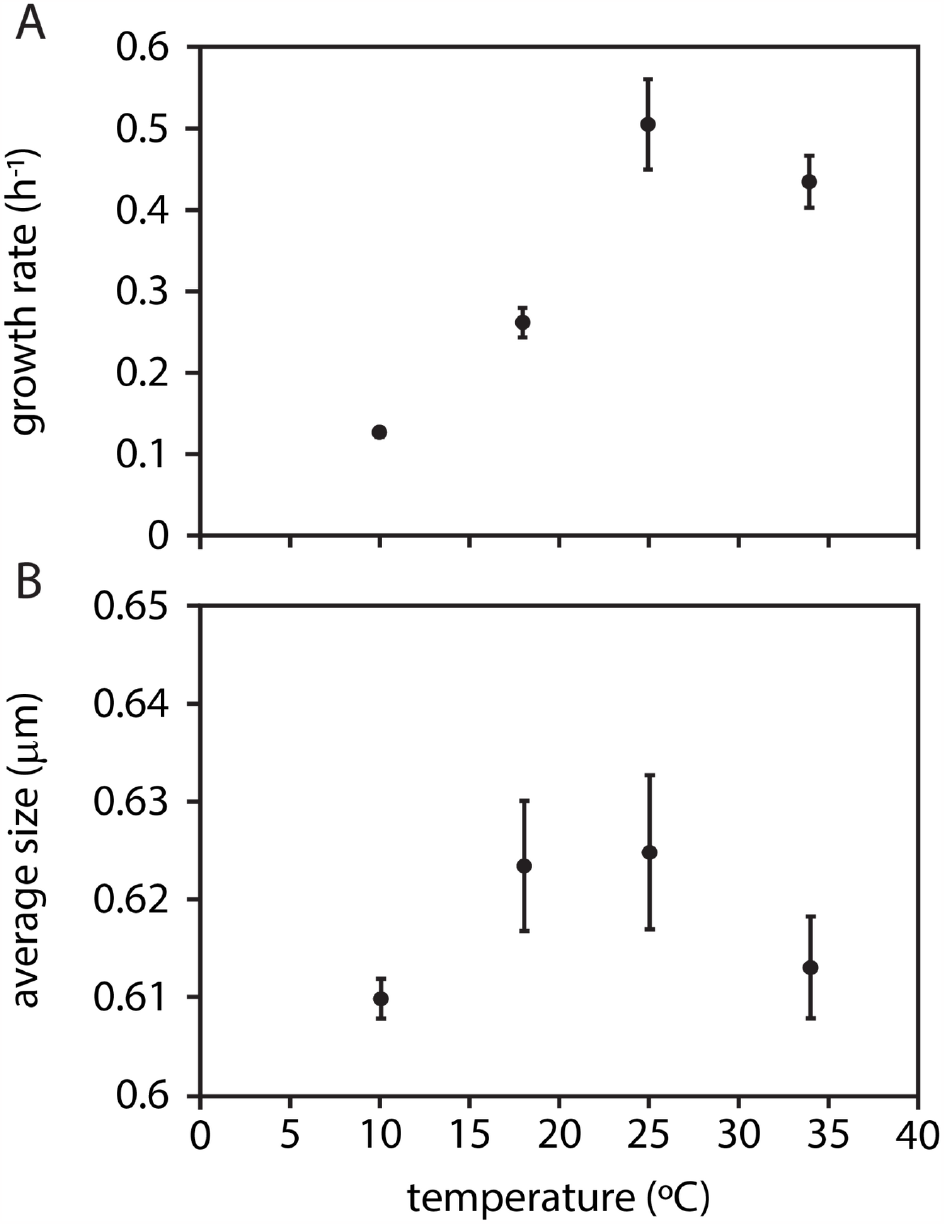
Growth rates and cell size in samples of *Dokdonia* sp. MED134 acclimated at different temperature conditions. (A) Growth rate and (B) average cell diameter along the thermal gradient under experimental conditions.

### Analysis of changes in gene expression under light and dark conditions across the thermal gradient

Following the acclimation period at each temperature, RNA samples for transcriptomic analysis were collected during exponential growth under light and dark conditions (i.e., after 2 hours of dark exposure; see Materials and Methods). First, an analysis of differential expression was performed between samples collected under light and dark conditions for all temperature treatments by DeSeq2. Employing a fold-change cutoff value of 0.5, only 15 DE genes were found by light conditions in all four temperature values (Table S2), which represents less than 0.5% of MED134 gene set. This result indicates that the impact of light versus dark conditions was only mild on the global gene expression patterns of MED134 under our experimental setting, which included day/night cycles. Considering the individual temperatures assessed, we found that the highest number of DE ORFs between light and dark conditions was found at 18ºC (41 genes overexpressed in the light and 3 in the dark), followed by 10ºC (32 genes overexpressed in the light and 2 in the dark), 34ºC (31 genes in the light and none in the dark) and 25ºC (20 in the light and 8 overexpressed in the dark, Table S2, Figure S1).

Only one gene was significantly upregulated under dark conditions at all temperatures except at 34ºC (Wald test, α = 0.01), encoding a regulatory two-component system sensor His kinase with a GAF domain (MED134_06219, Figure 2). The remaining DE genes identified were consistently upregulated under light conditions and were related to light sensing, light-induced cellular damage repair, and phototrophy (Table S2). The latter category included the PR (MED134_07119), and three PR-related genes involved in the synthesis of retinal molecules attached to the PR apoprotein (phytoene synthase and phytoene dehydrogenase; *crtBI*) or with a regulatory function (transcriptional regulator gene of MerR family, Figure 2). Other ORFs upregulated by light were a deoxyribodipyrimidine photo-lyase class I (MED134_14266), a regulatory gene activated by blue light (MED134_07089), an enzyme to repair damaged proteins (Met sulfoxide reductase; MED134_10226) and three genes predicted to be part of the same operon: a cryptochrome of the DASH family, a deoxyribodipyrimidine photolyase-related protein and a DNA photolyase/cryptochrome (MED134_10201, MED134_10206, and MED134_10211, respectively, Figure 2).

**Figure 2.**
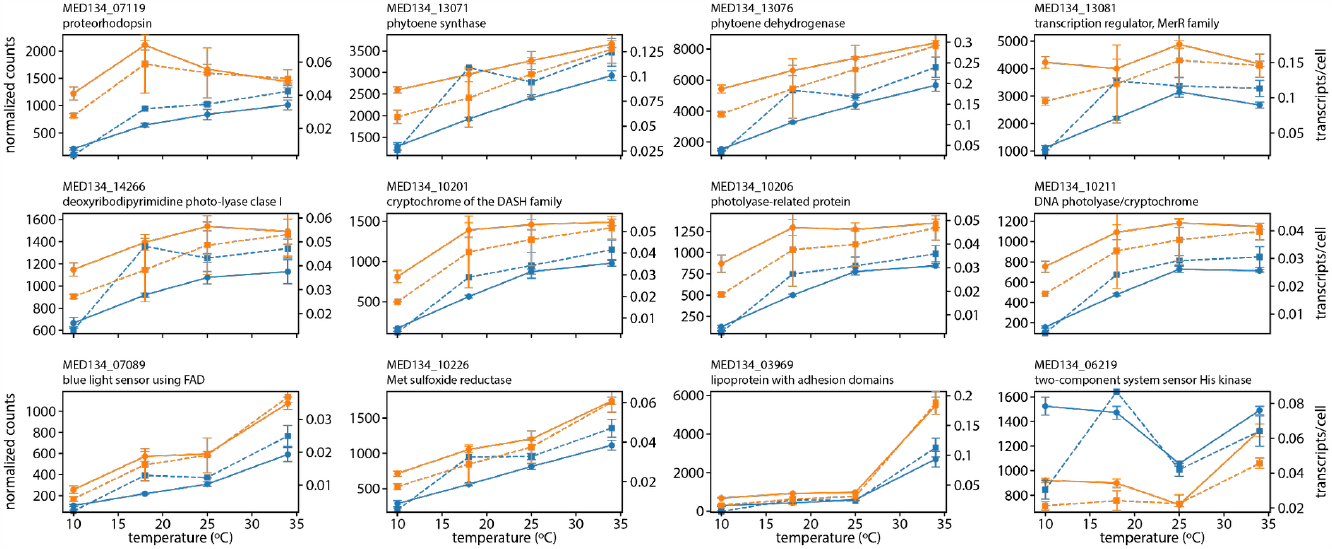
Gene expression values of key genes in light and dark conditions across various temperature treatments. All genes, with the exception of MED134_10226 and MED134_03969, were differentially expressed across all temperatures (Wald test, α = 0.01). Solid lines correspond to mean DeSeq2-normalized counts across all samples at the same temperature, and error bars to the standard error of the mean. Dashed lines represent average transcript abundances (transcript/cell values). Data corresponding to light treatment is depicted in orange, whereas blue represents dark.

In general, the patterns of expression of the latter genes along the thermal gradient were highly consistent in terms of DeSeq2-normalized values and estimates of transcript per cell (Figure 2). Genes upregulated by light generally showed minimum values of expression at the cold threshold of growth (10ºC). This was the case for the PR, which additionally exhibited minimum differences in the ratios of light/dark expression at the warmest temperature. The light/dark fold-change expression ratio of the PR gene was 2.6 at 10ºC, 1.7 at 18ºC, and 1 at 25ºC, while there was no differential expression at 34ºC (DeSeq2-normalized values). By contrast, a predicted extracellular lipoprotein with adhesion domains (MED134_03969) showed the maximum light/dark fold-change expression ratio at the warm threshold of growth (Figure 2).

### Analysis of gene expression patterns along the thermal niche

In contrast to the limited number of genes DE by light conditions, a high proportion of MED134 genes (1930 genes, 67% of MED134 genes) were DE across temperatures (LTR test, α = 0.01; see Materials and Methods). Considering only DE genes, clusters with similar expression patterns along the thermal niche were identified by using *Clust* and DeSeq2-transformed gene counts as input (Figure 3). All replicates were pooled per temperature irrespective of the light/dark condition and genes that were DE by light were excluded from the analysis. Five clusters were identified by *Clust*, including a total of 1204 genes (42% of the MED134 gene set, Table S3). These clusters showed two main transcriptional responses: either a positive response with temperature (Clusters 0, 2, and 3) or a negative response (Clusters 1 and 4; Figure 3). In the clusters of the first category, the upregulation trend was observed over the whole temperature range (Cluster 0), only under warm stress conditions (i.e. 34ºC; Cluster 2), or following the pattern of growth rates (with an upregulation by temperature only over the range 10 to 25ºC; Cluster 3). Clusters 1 and 4 showed the inverse response pattern to those of Clusters 0 and 3, respectively, with either a downregulation over the whole temperature range or only between 10 and 25ºC (Figure 3).

**Figure 3.**
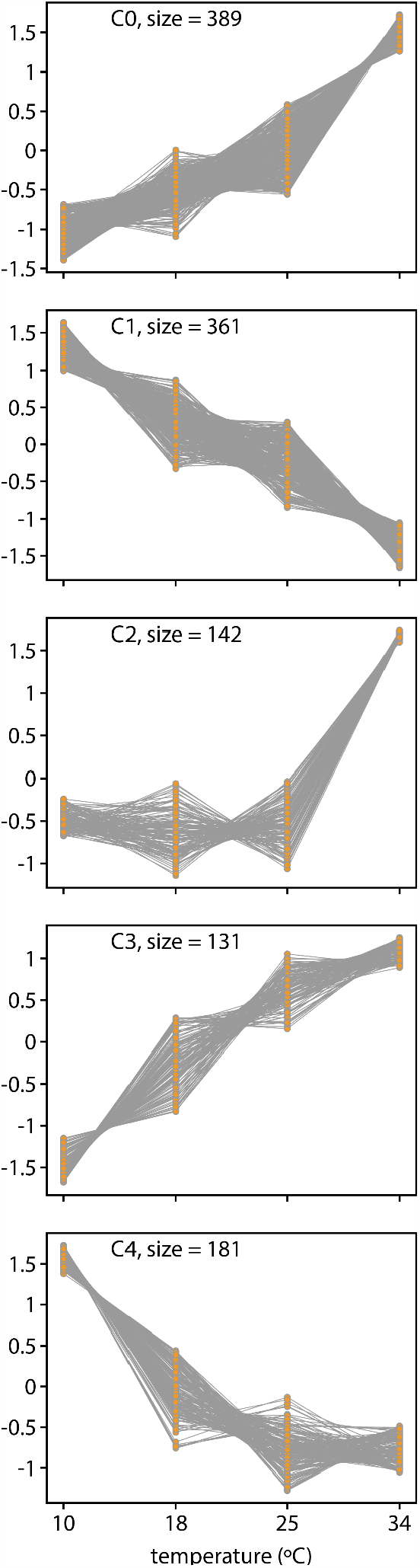
Identified clusters of across-temperature gene expression patterns. Clusters were constructed using a z-score normalization of the DeSeq2-transformed counts (values shown here). Genes that were differentially expressed between light and dark conditions (Wald test, ɑ = 0.01) were removed and light and dark samples were merged before clustering.

Over-represented metabolic pathways in each cluster were identified with *clusterProfiler*. However, on average, 78% of the genes in each cluster were not functionally annotated in KEGG, which limited the analysis. Over-represented pathways were only found for the two clusters which showed a negative response with temperature; while the subsystem Ribosome was over-represented in Cluster 1, the pathways Arginine biosynthesis, Biosynthesis of amino acids, Alanine, aspartate and glutamate metabolism, and Biosynthesis of secondary metabolites were over-represented in Cluster 4 (Table S4). The cold upregulation of some individual marker genes in these pathways can be visualized in Figure 4, where estimates of transcript abundance per cell have been included along with DeSeq2-normalized values.

**Figure 4.**
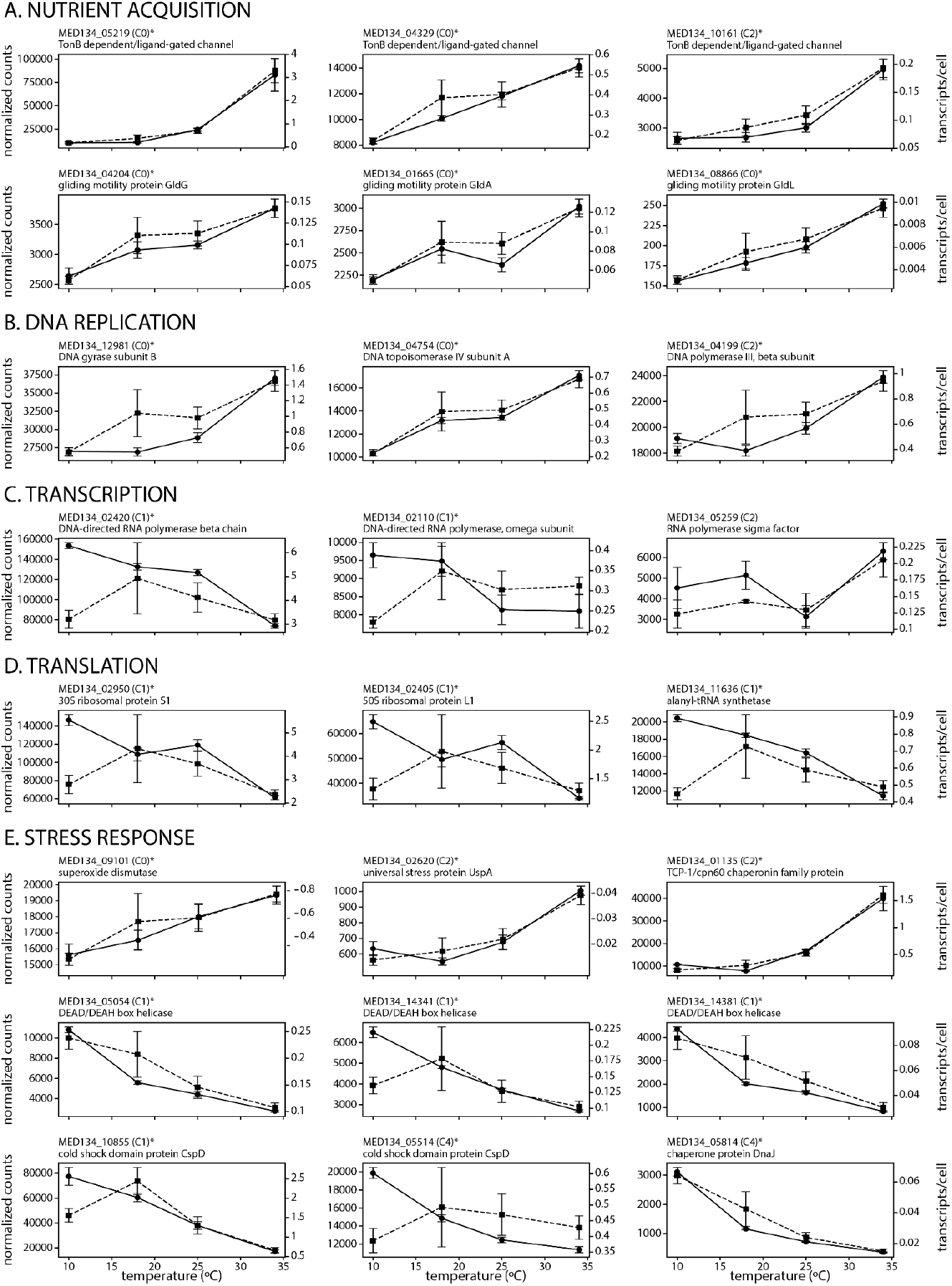
Expression patterns across the temperature range for selected key functional genes. Estimates of transcript abundance per cell (dashed lines) and Deseq2-normalized values (solid lines) are displayed along the temperature range. In both cases, values represent averages across replicates, with error bars corresponding to the standard error of the mean. Locus tags as well as the assigned cluster (using DeSeq2-normalized values) are shown in each case. Asterisks denote differential expression along the temperature gradient (LTR test, ɑ = 0.01).

While similar trends were generally found by both approaches over the range of 18ºC to 34ºC, contrasting results were found at 10ºC in genes assigned to Clusters 1 and 4, with an upregulation at the cold threshold of growth only observed in DeSeq2-normalized values. This pattern was also clear in highly-expressed marker genes involved in cellular transcription (i.e. the RNA polymerase beta chain, Figure 4C) and translation (ribosomal proteins and aminoacyl-tRNA genes, Figure 4D). Such contrasting results between both approaches were likely due to the marked reduction in the global abundance of transcripts per cell observed at 10ºC, with median expression values 46% lower than the median average across temperatures, resulting in a downregulation of most genes at the latter temperature.

Only 10 genes showed a distinct decreasing pattern from 10ºC to 34ºC when looking at transcript per cell estimates, and all of them were included in clusters C0 and C4 when using DeSeq2-normalized values (Figure S2). These included genes involved in the stabilization of the secondary structure of proteins and nucleic acids, such as DEAD/DEAH box helicases (MED134_05054, MED134_14381, MED134_05809). A clear cold upregulation of other genes involved in the stress response, with a role as cellular chaperones such as *dnaJ* (MED134_05814) and the cold shock protein domain *cspD* (Figure 4E), was also observed both in DeSeq2-normalized values and estimates of transcript abundance. Conversely, other stress-related genes were upregulated toward warm conditions, such as those encoding the universal stress protein UspA and the superoxide dismutase (Figure 4E).

Among the highly-expressed genes that showed a clear upregulation towards warm conditions, we found some TonB transporters (TBDT) involved in taking up nutrients to the periplasm of the cell, gliding motility genes (*gldG, glgA, glgL*, Figure 4A) and genes related to DNA replication (Figure 4B). The latter genes included DNA gyrase subunit B, DNA polymerase III subunit β, and the DNA topoisomerase IV subunit A. For all these genes, the estimates of Deseq2 and transcript abundance per cell were highly consistent (Figure 4A, B). Notably, the expression of other highly-expressed genes followed the pattern of growth rates along the thermal niche (Cluster 3), including those encoding the TonB transporter with the largest median expression value (MED134_11471), the Na^+^/H^+^ antiporter NhaC (MED134_03169) likely involved in maintaining Na^+^ homeostasis, a zinc carboxypeptidase (MED134_03030), a metal-dependent amidohydrolase (MED134_13906), and a glycosyl transferase of type group 1 (MED134_05959) (Table S3). In Cluster 3, we also found the subunit ɑ of the acetyl-CoA carboxylase carboxyl transferase (MED134_01520), involved in fatty acid synthesis, and several genes involved in anaplerotic carbon fixation, such as the PEP carboxylase (MED134_06244) and the PEP carboxykinase (MED134_10331, Figure 5). In the TCA cycle, most genes were similarly expressed in terms of transcript abundance, at least in the range 18ºC to 34ºC, with some notable exceptions: the gene encoding the isocitrate dehydrogenase, responsible for the dehydrogenation of the isocitrate to 2-oxoglutarate (MED134_14141) was upregulated towards warm conditions, while marker genes of the glyoxylate shunt (i.e., isocitrate lyase and malate synthase) were only clearly upregulated at the warmest threshold of growth (Figure 5).

**Figure 5.**
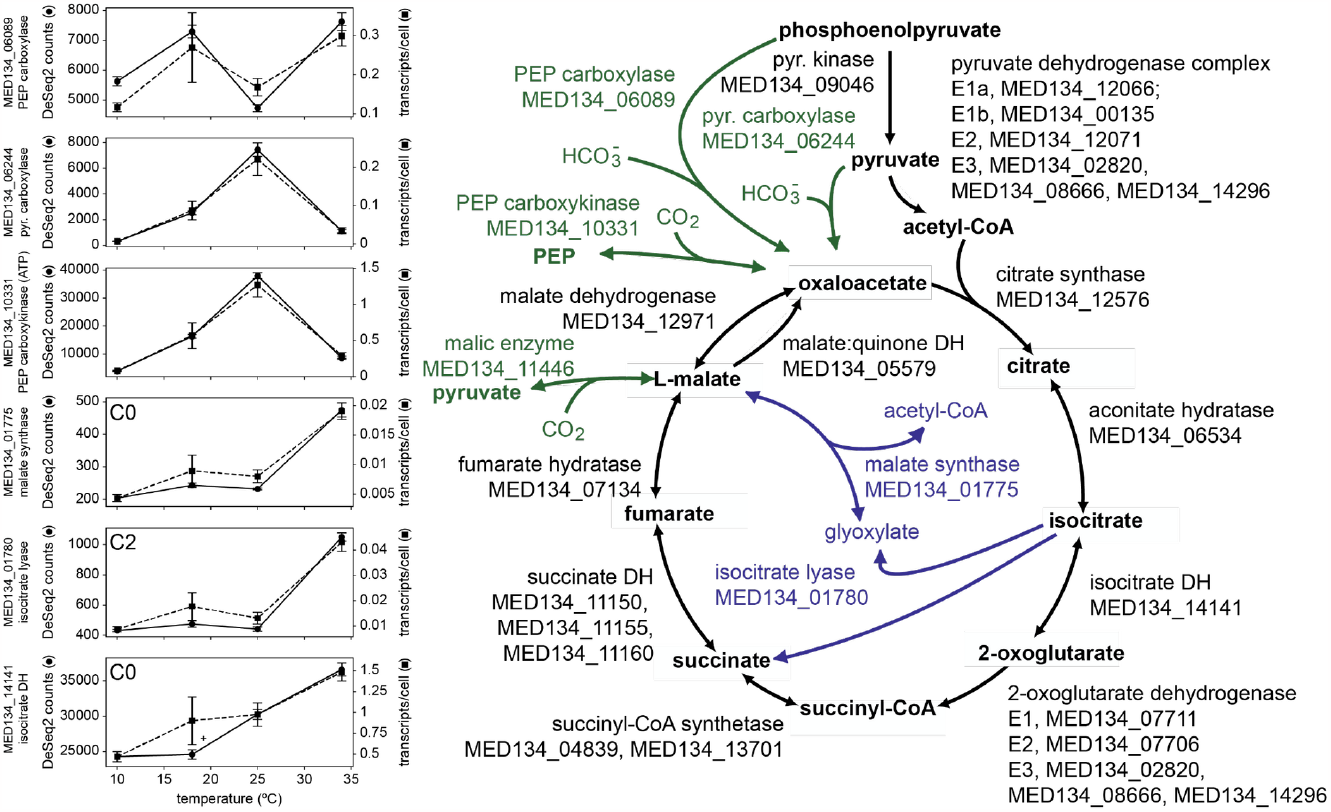
Expression patterns across the temperature range for genes belonging to the TCA cycle and glyoxylate shunt in *Dokdonia* MED134. Estimates of transcript abundance per cell (dashed lines) and Deseq2-normalized values (solid lines) are displayed along the temperature range. All genes, except for genes in the glyoxylate shunt (highlighted in blue) and anaplerotic enzymes (in green), were DE across temperatures (LTR test, ɑ = 0.01). Locus tags as well as the assigned cluster (using DeSeq2-normalized values) are shown in each case. DH indicates dehydrogenase; PEP, phosphoenolpyruvate, and pyr stands for pyruvate. The rest of DE genes in the figure are shown in Figure S3.

Finally, similar to most genes in the TCA cycle, a large representation of highly-expressed genes participating in housekeeping functions such as glycolysis and gluconeogenesis, oxidative phosphorylation, or nucleotide and amino acid metabolism were not DE by temperature (LTR test, ɑ = 0.01, a total of 860). The same pattern was found for highly conserved bacterial genes (i.e., RNA polymerase subunit genes and transcription factors) and some of the most highly-expressed genes which are part of operons containing a TDBT (e.g., MED134_12371-MED134_12381 and MED134_10780-MED134_10820, Table S3). TBDT and lipoprotein genes might be equivalent to the SusCD system in *Bacteroides* species, involved in the binding and uptake of a wide range of polysaccharides from the environment (Kuwahara et al. 2004; Xu et al. 2003). Outer membrane proteins (MED134_10815, MED134_07606, MED134_09311) or a fungalysin metallopeptidase type M36 with the *Por* domain, were also highly expressed under all temperature conditions.

## DISCUSSION

Light on the surface of the ocean is a plentiful source of energy for ca. half of marine bacterioplankton cells, which contain light-activated energy-generating rhodopsins (Venter et al. 2004). This feature is prominent in marine flavobacteria, which also dominate the expression of PR seasonally (Arandia-Gorostidi et al. 2020). Phototrophy, far from being an ancillary adaptation to the conditions in the ocean, seems to be an important trait for this group. The genus *Dokdonia*, and particularly our model strain MED134, has been one of the few model organisms where a stimulatory effect of light on growth has been shown experimentally (Gómez-Consarnau et al. 2016; 2007; Kimura et al. 2011). The molecular mechanisms that make *Dokdonia* benefit from light for growth are not fully understood.

However, previous transcriptomic analyses have shown that light might induce a remodeling of the carbon flow inside the cell through the glyoxylate shunt (Palovaara et al. 2014). We did not observe a significant effect of light on the abundance of transcripts of the glyoxylate shunt under our experimental conditions. This is likely related to the fact that, by contrast to previous studies where *Dokdonia* MED134 was grown under continuous light or dark conditions (Palovaara et al. 2014; Gómez-Consarnau et al. 2016), we grew this strain under light/dark cycles to simulate natural conditions in typical temperate systems. Therefore, our results indicate that some of the transcriptional differences previously reported under continuous light *versus* dark conditions are likely attenuated under daily light cycles.

When growing under diel cycles, light-protecting mechanisms were upregulated during the light period in MED134, such as photolyases and cryptochromes, which might repair damaged DNA (Essen 2006), act as blue light sensors (Sancar 2003), or even be involved in both activities (Bayram et al. 2008). This suggests that visible light, besides representing an extra energy source for MED134, may also induce cellular stress in this flavobacterium. PR, as a marker gene of the photoheterotrophic lifestyle in marine bacteria, was also clearly upregulated under light conditions, supporting the view that the light-induced expression of this gene is an inherent feature of PR-containing flavobacteria, both in culture (Gómez-Consarnau et al. 2016; Feng et al. 2013) and in the environment (Lami et al. 2009). Notably, in MED134, light upregulated the expression of PR at all temperatures except at 34ºC, indicating a possible limitation in the regulation of this key gene under warm stress conditions.

It has been suggested that the H^+^ gradient supported by the PR activity is used to power TBDT systems (Morris et al. 2010). These systems can be tailored to the uptake of peptides, which are readily consumed by flavobacteria for growth (Liu, Wawrik, and Liu 2017; Simon et al. 2012; Cottrell and Kirchman 2000) and are part of the Sus complex, which mediates the binding of complex substrates to the cell surface to perform a partial degradation before incorporating the products to the periplasm (Anderson and Salyers 1989a; 1989b; Kuwahara et al. 2004; Xu et al. 2003). We found that genes encoding TBDT systems were among the most actively expressed in MED134, in consistency with previous results for the same strain (Gómez-Consarnau et al. 2016) and, more generally, in environmental *Flavobacteriia* (Arandia-Gorostidi et al. 2020). While some TBDT genes were constitutively expressed under different light and temperature conditions, others were upregulated towards warm conditions, similar to some genes involved in gliding motility (Figure 4A). This type of motility is also a distinctive feature in *Bacteroidetes*, which helps the bacterium slide over surfaces, such as particles, in search of nutrients (M. J. McBride 2001; Mark J. McBride 2004). Thus, our results suggest that the uptake of selective carbon and nutrient sources via the activation of gliding motion and specific TBDT may be enhanced to promote cell metabolism under warm temperature conditions. Accordingly, a sharp decrease in gliding motility was observed in a flavobacterium strain (*Cytophaga* sp. U67) acclimated to 25ºC after a shift to 13ºC (McGrath, Moss, and Burchard 1990). Increased metabolic rates under warm conditions may also induce a higher oxidative stress, which is reflected in the upregulation of the superoxide dismutase under warm conditions. Conversely, cold conditions upregulated the expression of DEAD-BOX helicases and chaperones, which are components of the universal cold-shock response (Weber and Marahiel 2003).

In terms of carbon metabolism, we found that the expression of major pathways such as glycolysis, gluconeogenesis, and the TCA cycle was not substantially impacted by temperature, reinforcing the view that these are housekeeping genes for MED134. By contrast, the expression of marker genes involved in anaplerotic CO_2_ fixation and the glyoxylate shunt were correlated with growth rates over the thermal niche (i.e., PEP carboxylase and PEP carboxykinase) or activated at the warmest threshold of growth, respectively. Anaplerotic carbon fixation can represent up to 30% of total biomass production in this flavobacterium when growing on alanine (Palovaara et al. 2014), which is the carbon source that we used in our experiments. Thus, our results agree with the idea that anaplerotic metabolism has an important role in biomass production in MED134, maximizing the expression of this pathway at the optimum growth temperature. On the other hand, the upregulation of the glyoxylate shunt at 34ºC may reflect a preferential growth dependence on C_2_ compounds under warm stress conditions, bypassing the two TCA cycle steps where CO_2_ molecules are released. Our results, together with previous results with the same strain MED132 (Palovaara et al. 2014), indicate that the expression of the glyoxylate pathway is rather sensitive to changes in growth conditions. Clearly, a more in-depth understanding of the relevance of anaplerotic C fixation and the glyoxylate shunt for the carbon metabolism of marine photoheterotrophs under different environmental conditions is needed.

In summary, our results indicate that the model photoheterotroph MED134 can sense and respond to light variations over the daily cycle. However, their transcriptional response to light was relatively mild as compared to the gene expression changes induced by temperature variations within their thermal tolerance range. Cold temperature upregulated genes of the canonical cold-shock response as well as pathways related to cellular translation, likely to compensate for a decreased efficiency of the protein synthesis machinery at low temperatures (Toseland et al. 2013). By contrast, the expression of most genes involved in the TCA cycle, glycolysis, or gluconeogenesis was not differentially expressed along the thermal niche, suggesting that the transcriptional regulation of these major carbon metabolism pathways was resilient to temperature stress. However, anaplerotic C fixation and the glyoxylate shunt were clearly impacted by temperature conditions, supporting the idea that these key pathways may induce changes in carbon remodeling in *Dokdonia* depending on growth conditions. Finally, we also found a significant response to temperature for some highly-expressed genes likely involved in the uptake and degradation of complex carbon compounds, as well as in light energy transduction, showing that our model Bacteroidetes organism gradually adapts the expression of both sets of genes, likely to compensate for suboptimal growth conditions.

## ACKNOWLEDGEMENTS

We are grateful to the Spanish Ministry of Economy and Competitiveness (MINECO) for supporting L.A.-S.’s Ramón y Cajal research contract (RYC-2012-11404), A.S.P.’s FPI Ph.D. fellowship (BES2015076149), and the project TECCAM (CTM2014-58564-R). JMG and SRE were supported by project PID2019-110011RB-C32 (Spanish Ministry of Science and Innovation, Spanish State Research Agency, doi: 10.13039/501100011033).

