## Supplementary material for "Gene expression response to temperature acclimation in a model photoheterotrophic marine flavobacterium (*Dokdonia* sp. MED134)": Figure S1

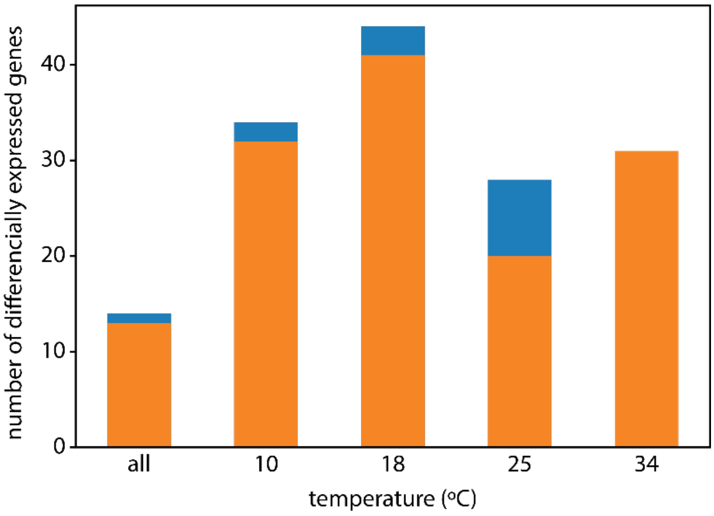


**Figure S1. Number of differentially expressed genes across temperature treatments.** Number of differentially expressed genes between light and dark conditions for all four temperature treatments (Wald test, α = 0.01, fold change cutoff = 0.5). Genes with higher expression levels in the light are depicted in orange while blue corresponds to dark.
