## Supplementary material for "Gene expression response to temperature acclimation in a model photoheterotrophic marine flavobacterium (*Dokdonia* sp. MED134)": Figure S2

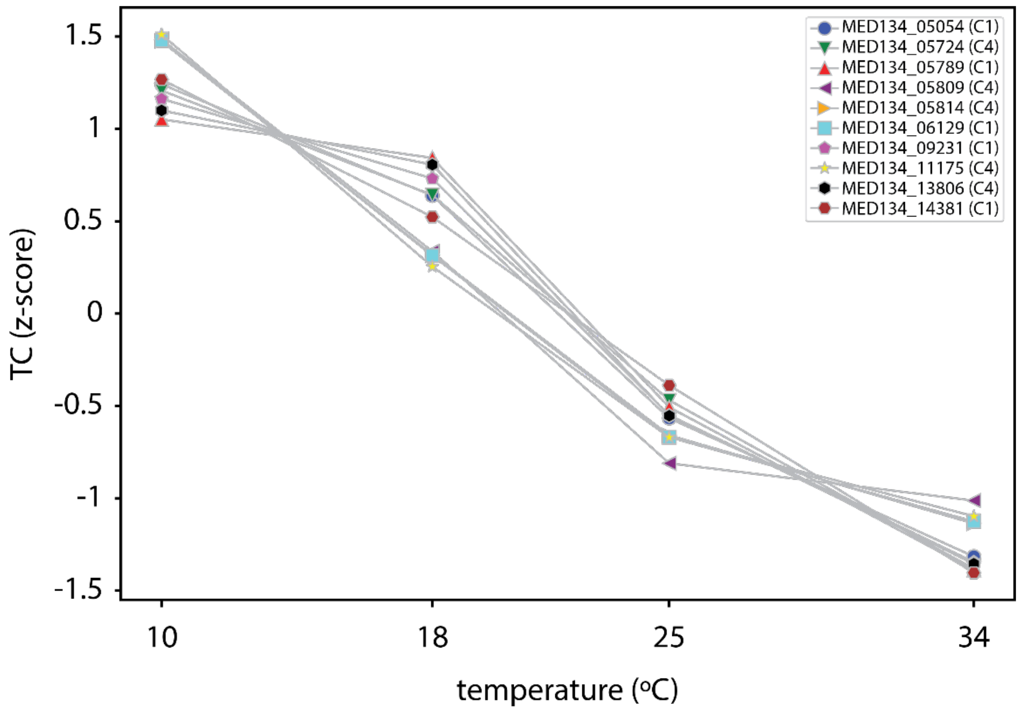


**Figure S2. Genes with strictly decreasing transcript per cell values for increasing temperatures.** A total of 10 genes in MED134 showed a strictly decreasing expression pattern when using transcript per cell estimates. All these genes were included in clusters C1 and C4 obtained with DeSeq2-normalized values. The cluster ID is shown within parentheses in the legend.
