## Supplementary material for "Gene expression response to temperature acclimation in a model photoheterotrophic marine flavobacterium (*Dokdonia* sp. MED134)": Figure S3

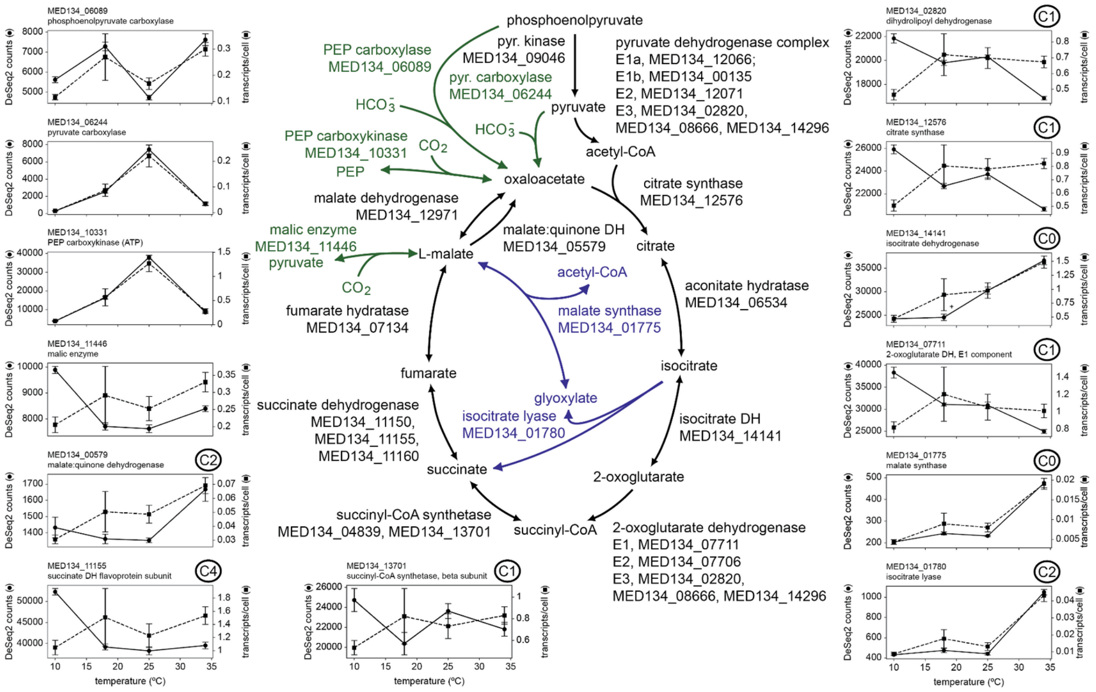


**Figure S3. Expression patterns across the temperature range for genes belonging to the TCA cycle and glyoxylate shunt in *Dokdonia* MED134.** Estimates of transcript abundance per cell (dashed lines) and Deseq2-normalized values (solid lines) are displayed along the temperature range**.** All genes, except for genes in the glyoxylate shunt (highlighted in blue) and anaplerotic enzymes (in green), were DE across temperatures (LTR test, α = 0.01). Locus tags as well as the assigned cluster (using DeSeq2-normalized values) are shown in each case. DH indicates dehydrogenase; PEP, phosphoenolpyruvate, and pyr stands for pyruvate.
